# Organization and priming of long-term memory representations with two-phase plasticity

**DOI:** 10.1101/2021.04.15.439982

**Authors:** Jannik Luboeinski, Christian Tetzlaff

## Abstract

Synaptic tagging and capture (STC) is a molecular mechanism that accounts for the consolidation of synaptic changes induced by plasticity. To link this mechanism to long-term memory and thereby to the level of behavior, its dynamics on the level of recurrent networks have to be understood. To this end, we employ a biologically detailed neural network model of spiking neurons featuring STC, which models the learning and consolidation of long-term memory representations. Using this model, we investigate the effects of different organizational paradigms of multiple memory representations, and demonstrate a proof of principle for priming on long timescales. We examine these effects considering the spontaneous activation of memory representations as the network is driven by background noise. Our first finding is that the order in which the memory representations are learned significantly biases the likelihood of spontaneous activation towards more recently learned memory representations. Secondly, we find that hub-like structures counter this learning order effect for representations with less overlaps. We show that long-term depression is the mechanism underlying these findings, and that intermediate consolidation in between learning the individual representations strongly alters the described effects. Finally, we employ STC to demonstrate the priming of a long-term memory representation on a timescale of minutes to hours. As shown by these findings, our model provides a mechanistic synaptic and neuronal basis for known behavioral effects.

## Introduction

Synaptic tagging and capture (STC) provides a well-established hypothesis of the molecular processes underlying synaptic consolidation, which has been investigated by a multitude of experimental and theoretical studies [1, 2, 3, 4, 5, 6, 7, 8, 9]. However, the dynamics resulting from the interplay between STC and recurrent network structures, which are characteristic for brain structures like the hippocampus and the neocortex [10, 11], and their relation to long-term memory still remain unclear.

Neuronal activity causes the increase or decrease of the transmission strength of synapses via long-term potentiation (LTP) and long-term depression (LTD). The early phase of these phenomena typically lasts for up to a few hours [12, 13]. To be maintained, the synaptic changes have to undergo a consolidation process, which is described by the STC hypothesis [1, 4]. When a group of neurons in a recurrent neural network experiences strong (learning) stimulation, this causes the formation of a Hebbian cell assembly, which thence represents a memory of the stimulation. Hebbian cell assemblies are groups of neurons exhibiting particularly strong synaptic connections, giving rise to facilitated activation of the assembly neurons [14, 15, 16, 17]. In our previous study [9] we developed a computational model and showed that a memory representation is first encoded by the early-phase of long-term plasticity, while STC subsequently transfers the synaptic changes to the late phase, yielding a long-term memory representation. Here we develop this model further and investigate the link between STC, properties of *multiple* long-term memories and their behavioral implication.

In the brain, spontaneous activity is ubiquitous and has been shown to inherently arise from deterministic dynamics in neural networks [18, 19]. In theoretical models spontaneous activity is usually elicited by background noise. This approach is biologically well-grounded through matching power spectra of in vivo neuronal activity on the one hand and of the stochastic processes used to model background noise on the other hand [20, 21]. Spontaneous activity causes the reactivation of existent cell assemblies via neuronal avalanches [22, 23] and thereby becomes relevant for cognition. In the current study, we use the concept of reactivation by spontaneous activity to link the reactivation dynamics of multiple long-term memories organized by STC to behavioral experiments. Namely, we compare our modeling results to two different classes of experiments from cognitive psychology: free recall experiments, and priming experiments.

Free recall is a memory test that requires subjects to reproduce information from memory, for example a sequence of words, without cuing. While several models have been developed to describe free recall [24, 25, 26, 27, 28, 29], its biological underpinnings remain elusive. With respect to the serial position of the input items, free recall experiments typically exhibit three prominent characteristics [30, 31, 32, 33, 34]: 1. primacy, the better recall of the items learned first, 2. contiguity, the increased probability of recalling neighboring items, and 3. recency, the better recall of the most recently learned items. Primacy is commonly related to extended rehearsal, in which the recency effect could play an indirect role [35, 36, 37]. Recency was first thought to be a solely short-term memory-dependent effect, but has then also been found to persist in long-term memory [32, 36, 38, 39]. The recency effect is usually much stronger than the primacy effect [31, 32]. These effects are often referred to together as serial-position effect. The serial-position effect is found to be expressed differently in experiments with different delay times before recall [32, 40, 41]. Given these diverse findings, free recall experiments have essentially contributed to distinguish between a short-term and a long-term memory storage in the human brain [33]. Two recent studies [27, 28] have shown that long-term associations between items account for free recall rather than short-term associations as in previous models [24, 26], but there is also evidence that familiarity might not essentially influence free recall [42]. With our model, we link neuronal and synaptic dynamics including STC to emergent network principles to provide a mechanistic explanation of the recency effect in long-term memory representations.

Priming (not to be confused with primacy) describes a large set of psychological phenomena across a multitude of timescales, which can occur either consciously or unconsciously [43, 44, 45]. Priming phenomena have in common that the recall of a certain memory is enhanced, following its reactivation by a “priming stimulus”. Studies on priming often target processes in the intersection between psychology and economics, for instance, bias in deciding to buy a certain product. As an example, if we see an advertisement of some brand of orange juice that we have had before, and soon afterwards go to the supermarket to do the weekly shopping, the likelihood of buying exactly that brand seen in the advertisement will be increased. Here, the advertisement acts as a priming stimulus [43, 46]. For priming on a timescale of seconds, Mongillo et al. [47] provided a model featuring the interplay of short-term plasticity and long-term plasticity. The authors also discussed the possibility of transferring this concept to the processes on the next, longer timescales: early- and late-phase long-term plasticity. We take on this idea and show that the STC mechanisms can account for a priming effect on a timescale of hours.

## Results

In this study, we employ a biologically plausible spiking recurrent neural network model, based on a previous model [9] of memory consolidation via synaptic tagging and capture (STC). In a proof-of-principle approach, we investigate multiple paradigms of the formation, consolidation, and interaction of three cell assemblies in the network. Measuring the likelihood of activation of these three assemblies, we show that two factors mainly guide their activation: 1. the learning order and 2. the type of overlaps between the assemblies. By mechanistically describing the learning order effect, our model provides a possible neural correlate for the recency effect in free recall experiments. Furthermore, we encounter differential effects for hub-like overlap structures, indicating that our model may provide neural correlates for the diverse behavior observed for similarity of memories, such as retroactive interference and enhanced recall [48, 49, 50, 51, 52]. A major focus of our study is on the role of consolidation via STC. We show that the emergence of the learning order effect and the effects in hub-like structures are strongly biased by a protocol with altered consolidation phases. Thereby, we make predictions on the memory dynamics on different time scales, which can be tested in behavioral experiments. As a final step, we show that the interaction between early- and late-phase long-term plasticity provides a molecular mechanism for priming phenomena lasting from minutes to hours.

The neural network that we use consists of 2500 excitatory and 625 inhibitory leaky integrate-and-fire neurons. To form a cell assembly, we apply strong stimulation to a subset of the excitatory neurons in the network and thereby elicit synaptic plasticity, which is modeled by a spike-timing-dependent calcium model and a firing rate condition. This early-phase plasticity is followed by consolidation via STC, which transfers weight changes to the long-lasting late phase. The synaptic and neuronal level of the model is depicted in Fig. 1a, linked to a schematic of the network level in Fig. 1b (for more details, see Methods and [9]).

**Figure 1:**
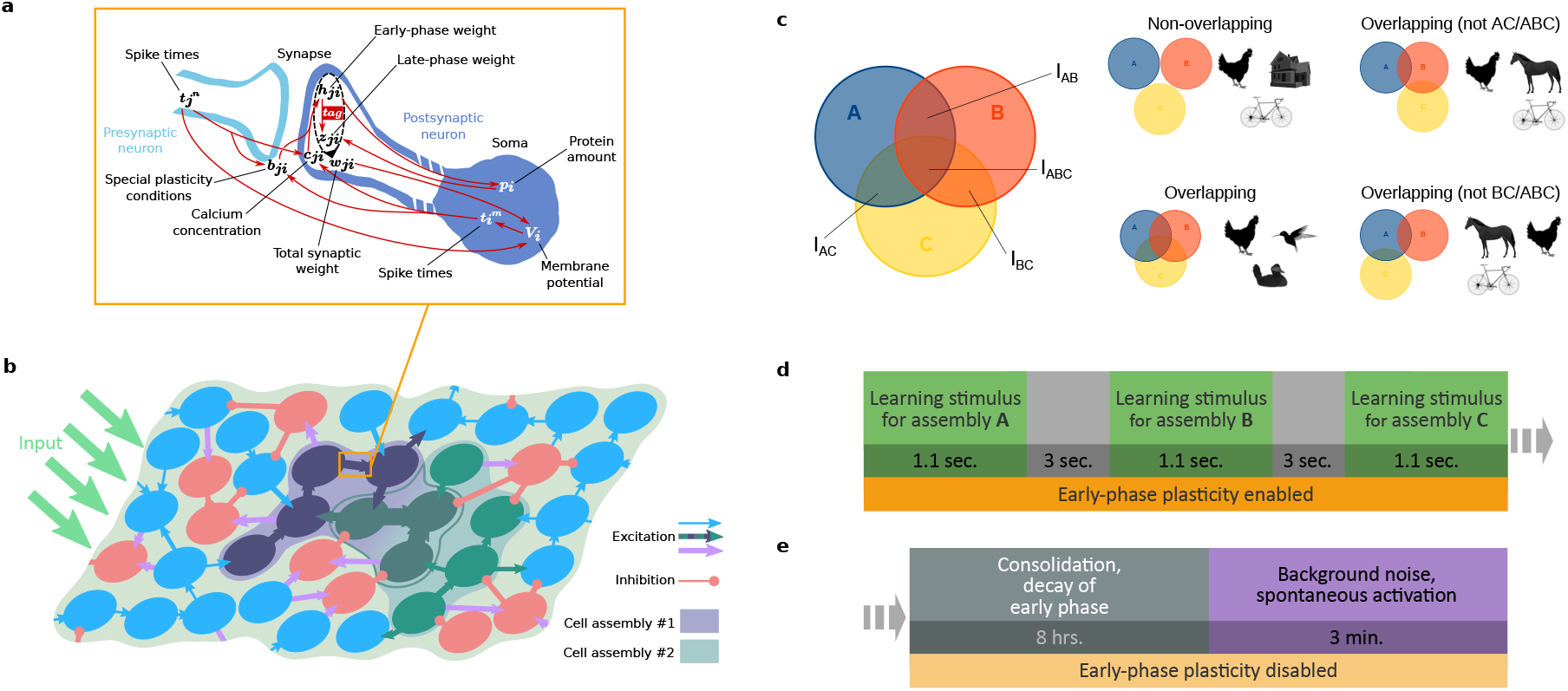
Model and considered protocols. **(a,b)** Schematic of the synaptic and network model (adapted from [9]). The synaptic model integrates the interplay between different mechanisms of calcium-dependent and rate-dependent synaptic plasticity and the STC hypothesis. The neural network consists of excitatory (light and dark blue and green disks) and inhibitory neurons (red disks) and receives external input from other brain areas. Only synapses between excitatory neurons undergo plastic changes via the processes shown in **a**. Hebbian cell assemblies representing memories consist of groups of strongly interconnected neurons (dark blue, green). The assemblies in **b** overlap by a fraction of their neurons (dark green). For more details see the main text. **(c)** Different organizational paradigms are defined by different overlap relationships between the three cell assemblies. Pictures show real-life examples (all adapted from pixabay.com). **(d)** Protocol of the learning phase: the three memory representations are learned sequentially, with three-seconds breaks in between. **(e)** Protocol for the consolidation and recall phase during which consolidation takes place for eight hours after learning. Then, spontaneous activity in the network, driven by background noise, is observed for three minutes. Time spans in **d** and **e** are similar to those in free recall experiments [31, 33, 40].

We investigated the activation of the assemblies by considering neuronal avalanches within the assemblies [22]. To consider an assembly being active, we utilized the border value of the 99% quantile for the number of spikes within a time window. That means, we considered the 1% of the time windows with most spikes (for more details, see Methods and Suppl. Fig. S10). Comparing the results for the likelihood of avalanche occurrence to the average firing rates in corresponding paradigms, we found similar behavior (Suppl. Fig. S5).

### Learning and consolidation of multiple long-term memory representations

Before investigating the spontaneous activation of long-term memory representations, we will demonstrate how the network learns and consolidates them. We start from an unbiased network in which all weights have the same value. Then, learning proceeds according to the protocol which is shown in Fig. 1d, followed by consolidation and the investigation of the spontaneous activity, shown in Fig. 1e. Substantially extended breaks between learning stimuli, such that consolidation already occurs in those phases, are considered in a later part of this study. In the main text, we consider four organizational paradigms, shown in Fig. 1c.

The differences between these paradigms are visualized by small pictures. The different paradigms will be examined in the next subsection, while we restrict ourselves to the non-overlapping case in this subsection.

As a first step, after learning and consolidating three non-overlapping memory representations, we tested whether all three assemblies served as functional memory representations. To this end, we examined the ability to recall the learned and consolidated assemblies by applying stimulation to 20% of the neurons of each assembly, subsequently (Fig. 2a,b). We found the response caused by this stimulation to be stronger and to last longer than the response to the same stimulation in a naive network before learning (the example for *A* is shown in Fig. 2a,b in the two leftmost panels), which confirms the functionality of the memory representations. Moreover, we observed avalanches within the assemblies during spontaneous activity after learning and consolidation (Fig. 2a,b). Note that the synchronicity in the activity is also increased after learning and consolidation (Fig. 2a). Since the average excitatory synaptic weight in our network after learning and consolidation is increased (103.8%, integrated over all weights in Fig. 2c), this can be explained by the findings of Brunel [53], stating that strengthening of excitatory connections in the network causes a shift towards a regime of synchronized regular activity. Thereby, we conclude that our network operates close to the phase transition, and such criticality is typical for the occurrence of avalanches [22, 23].

**Figure 2:**
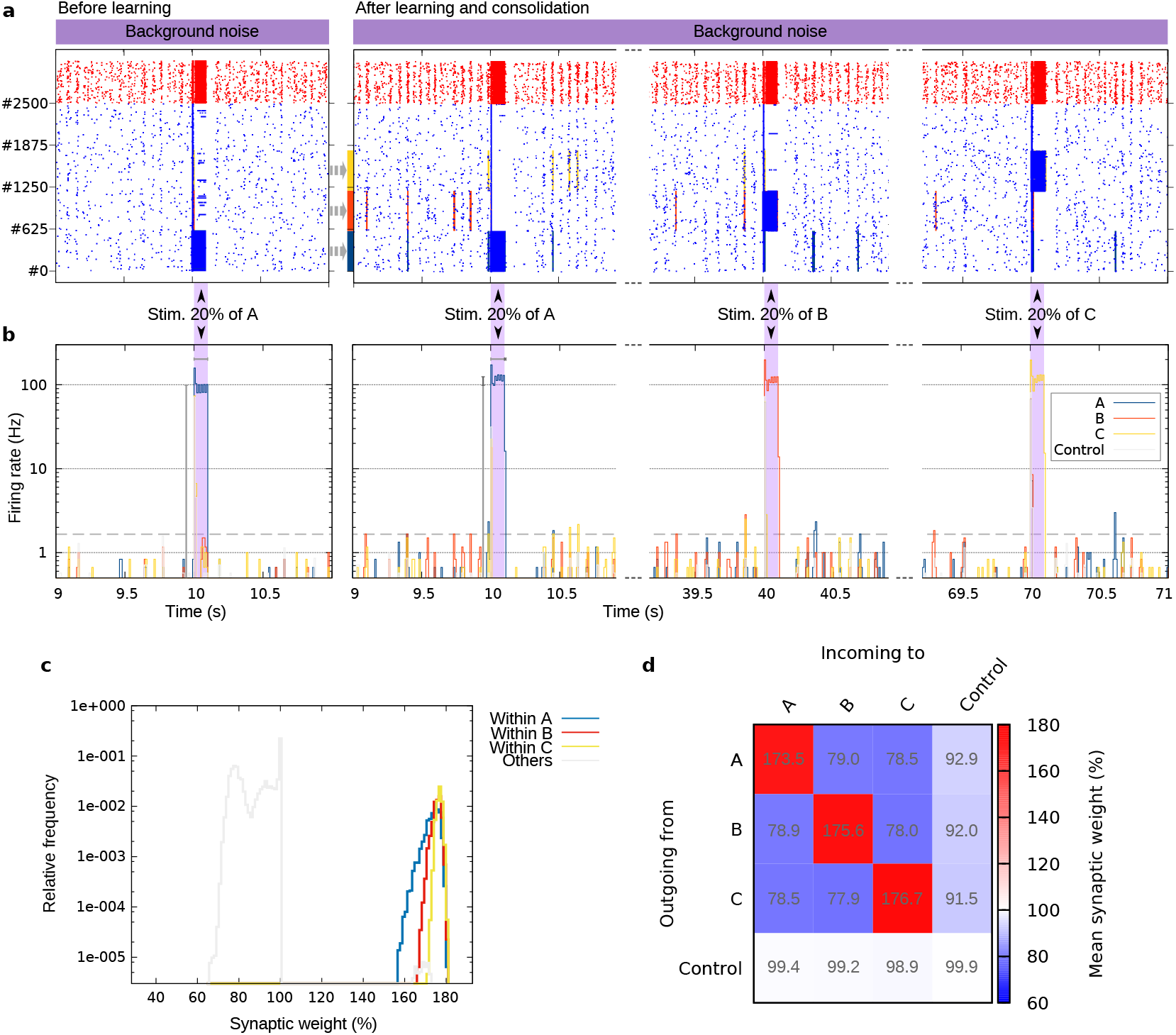
Three non-overlapping memory representations are successfully learned and consolidated. Learning order: *A*-*B*-*C*. **(a)** Raster plot showing the spiking in the whole network before and after learning and consolidation. At specified times, additional input is applied to 20% of the neurons of one assembly. Each assembly comprises 600 neurons. Spikes of excitatory and inhibitory neurons are shown by blue and red dots, respectively. Avalanches occurring in an assembly are indicated by bars in the respective color. **(b)** Firing rates corresponding to the raster plots (computed via bins of 10 ms). Shaded areas indicate the time frame of additional stimulation. The dashed line indicates the detection threshold for avalanches. In the two leftmost panels, gray bars next to the peaks emphasize the increase after consolidation in the mean value of the firing rate of *A* during the induced activation (+26.3%) and in the duration of the activation (+11.1%). **(c)** Weight distribution of the synapses within the assemblies, revealing the spread of LTP and LTD. **(d)** Abstract weight matrix showing the mean weight within and between all subpopulations of the excitatory population. The asymmetric nature of the matrix reveals the directionality of the coupling strengths. Data in **c** and **d** were averaged over 10 trials after learning and consolidation, and are relative to the initial value before learning.

In addition to the activity, we scrutinized the weight structure in the network after learning and consolidation. As expected, we found that the weights within each assembly are much higher than the weights in the remainder of the network (Fig. 2c). Remarkably, however, the weight distributions of the three assemblies differ. This is already a hint to the bias in assembly activation introduced by the learning order, which we will investigate in the following subsection. Complementary to the weight distribution, the abstract weight matrix in Fig. 2d reveals the directionality of the coupling strengths within and between subpopulations of the network after learning and consolidation. The matrix reflects the highly potentiated weights within each cell assembly, but also the depressed weights between the assemblies. Presumably, this depression arises from activation of the other assemblies upon learning a new assembly. The weights involving control neurons, on the other hand, exhibit only slight plasticity. Remarkably, synapses incoming from assembly neurons to control neurons are on average slightly depressed, while synapses outgoing from control neurons remain unchanged. Comparing with the weights after learning but before consolidation (Suppl. Fig. S1) reveals that at that time there is also depression in the synapses outgoing from control neurons, which is yet too weak to become consolidated. The depression of the weights presynaptic to the control neurons probably also emerges during learning of the assemblies, while the weights postsynaptic to the control neurons only receive indirect activation during learning. The higher calcium increase through presynaptic spikes in our model might increase this effect further.

For the overlapping paradigm, the weight distribution and the abstract weight matrix are given in Suppl. Figs. S2 and S3. Those results reveal further asymmetries for the weights of the intersections between assemblies, exemplifying the richness of our model.

### Learning order and overlaps guide spontaneous activity

After the learning and consolidation of the three assemblies, we systematically investigated the spontaneous reactivation of the assemblies driven by background noise. We collected data from three minutes, which is a time span similar to the recall period in free recall experiments [31, 33, 40]. A sample of spontaneous activity and the detected avalanches is shown in Fig. 3a,b.

**Figure 3:**
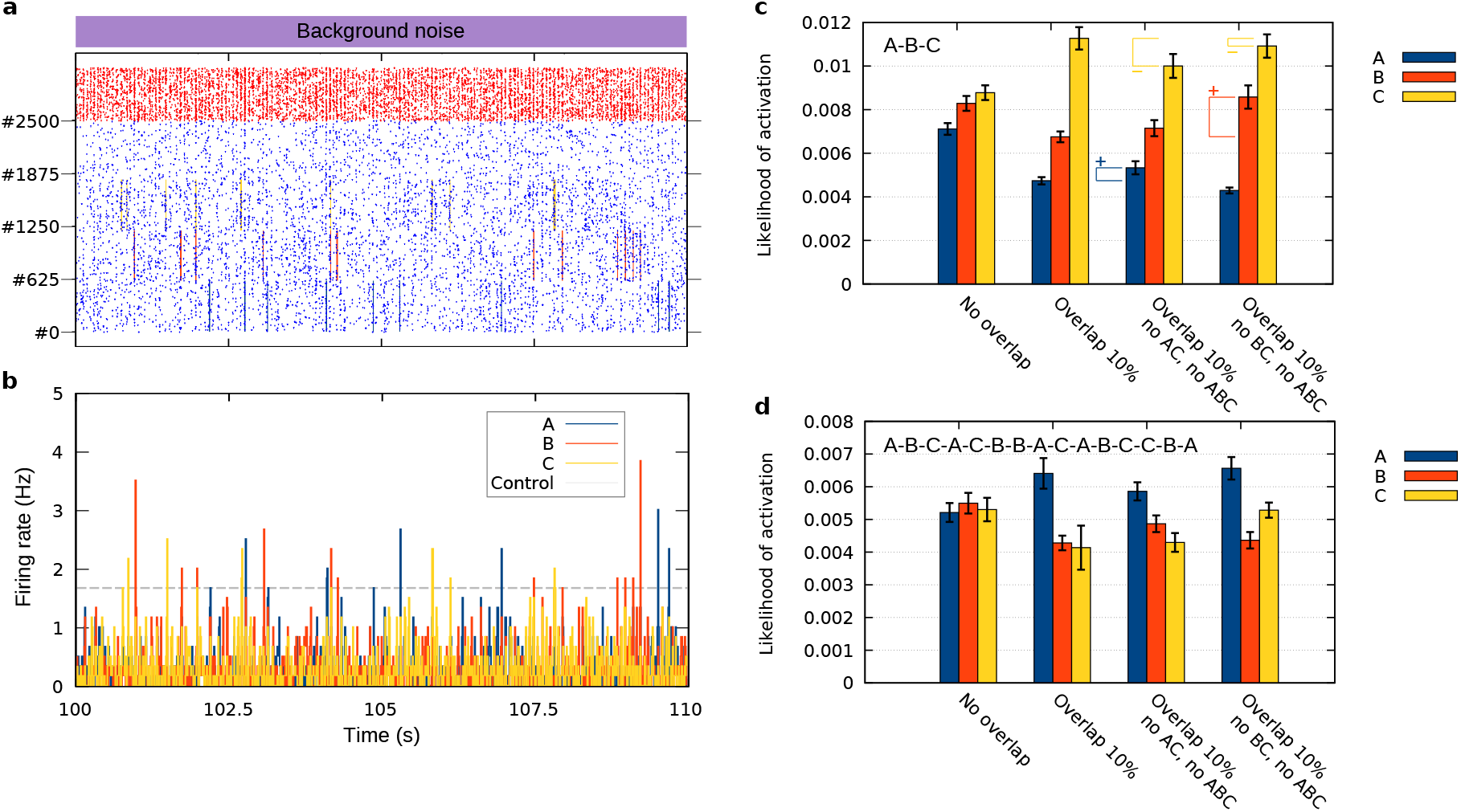
Spontaneous activity of long-term memory representations, driven by background noise. **(a)** Raster plot showing the spontaneous spiking activity in the whole network induced by background noise. Each assembly comprises 600 excitatory neurons (spikes indicated by blue dots). Spikes of inhibitory neurons are shown by red dots. Avalanches occurring in the assemblies are indicated by bars in the assembly colors. Assemblies do not overlap and are learned in the order: *A*-*B*-*C*. **(b)** Firing rates corresponding to the raster plot (computed via bins of 10 ms). The dashed line indicates the threshold for the detection of an avalanche. **(c)** Overview of the likelihood of avalanches in different organizational paradigms, learned with standard protocol (order: *A*-*B*-*C*). Averaged over mean likelihood from 10 networks. Error bars show the 95% confidence interval. Braces and plus/minus signs indicate differences to the “Overlap 10%” paradigm. **(d)** Overview of the likelihood of avalanches as in **c**, but learned with extended, interleaved protocol (order: *A*-*B*-*C*-*A*-*C*-*B*-*B*-*A*-*C*-*A*-*B*-*C*-*C*-*B*-*A*). Data in **c** and **d** were averaged over 10 trials.

Overlaps of memory representations on the network level are of huge relevance because they form the basis of memory similarity on the cognitive level [27, 29, 48, 51]. Here, we consider four different paradigms of overlaps, as they are shown in Fig. 1c (also see Methods). Small images picture real-life examples of non-overlapping and overlapping memory representations: representations of entities from three different categories, like chicken, house, and bicycle, do not overlap, while representations from the same category, like three types of birds, overlap. Mixed cases occur if two entities are in the same category while there is no common category for all three, for instance, bicycle and horse are both means of transportation, and chicken and horse are both farm animals. In the Supplementary Information to our study, we present results on more paradigms, including such with overlaps of varied percentage (Suppl. Figs. S5 and S6). For an overview over these paradigms, see Suppl. Table S1. The most important result from these additional paradigms is that in the majority of cases, larger overlaps further increase the effects that we describe in the following.

Considering the mean likelihood of activation in each of the four paradigms from Fig. 1c with assemblies that were learned in the order A-B-C, we encounter the first main result of our study: the learning order significantly biases the spontaneous activity, such that the activation of more recently learned assemblies is promoted (shown in Fig. 3c). While this is already the case without overlap, overlap of between assemblies increases the effect even further.

Our second finding is indicated in Fig. 3c by braces with a plus or minus sign. The braces demonstrate differences between the paradigms with less overlaps and the “Overlap 10%” paradigm. The lack of an overlap between two assemblies (in the abundance of the other overlaps) creates a hub-like structure, where the assembly that overlaps with the two other assemblies is the hub. We find that such hub-like structure reduces the learning order effect for the two assemblies with less overlaps. This means, the activation of the assembly that is disadvantaged by the learning order effect (*A* in the “no AC” paradigm, *B* in the “no BC” paradigm) is increased, while the activation of the assembly favored by the learning order effect (*C*) is decreased. As we will show below, this effect as well as the learning order effect itself are essentially caused by LTD.

For gathering further knowledge about the learning order effect and to examine other effects without the bias of the learning order, we tried to eradicate the learning order effect (also cf. the following subsection on priming). To this end, we employed a learning protocol for the interleaved learning of the three assemblies, using an extended sequence with the order *A*-*B*-*C*-*A*-*C*-*B*-*B*-*A*-*C*-*A*-*B*-*C*-*C*-*B*-*A* (random with the constraint that every assembly occurs equally often). Compared to Fig. 1d, the interleaved protocol simply contains more steps, but otherwise stays the same. In the non-overlapping paradigm, the interleaved learning successfully eradicates any significant impact of the learning order (Fig. 3d). In the other paradigms, however, a certain impact remains. Note that the most recently learned assembly in the extended sequence is *A*, which is also the mostly reactivated assembly.

In the next step, we considered a control case for the learning order *A*-*B*-*C* in which we blocked LTD, and found what had already been hinted by the distribution of weights resulting from the standard protocol (inset in Fig. 4a): the learning order effect vanishes if there is no LTD (Fig. 4a,c). Furthermore, the effect in hub-like structures that we discussed previously, vanishes as well. Instead, the activation of the hub assembly (*B* in the “no AC” paradigm, *A* in the “no BC” paradigm) clearly stands out, as it is expected for an LTP-only model. In addition, the overall likelihood of activation increases.

**Figure 4:**
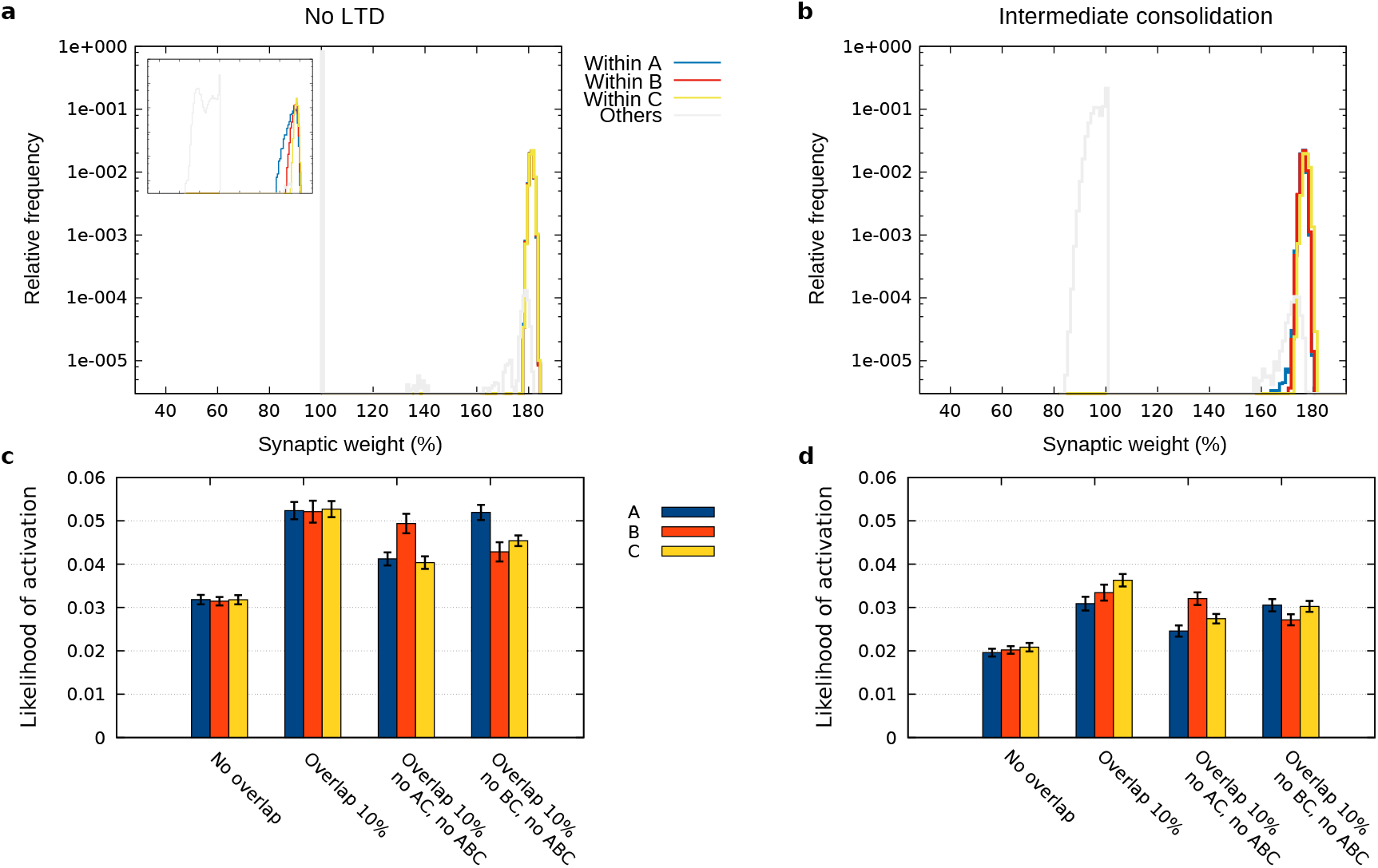
Scrutinizing the underpinnings of the learning order and overlap effects. **(a)** Weight distribution of the synapses within the assemblies, revealing that if LTD is blocked, the synapses belonging to the three assemblies are equally potentiated (non-overlapping case). The inset repeats the plot from Fig. 2c for comparison. **(b)** Weight distribution of the synapses within the assemblies learned with a modified standard protocol where the breaks between learning *A*, *B*, and *C* last 8 hours instead of 3 seconds (intermediate consolidation). The plot reveals a similar structure as in the no-LTD case. **(c)** Overview of the likelihood of avalanches in different organizational paradigms, learned with standard protocol (*A*-*B*-*C*), with LTD blocked. **(d)** Overview of the likelihood of avalanches in different organizational paradigms for the intermediate consolidation case (order *A*-*B*-*C*). Data were averaged over 10 trials. Error bars show the 95% confidence interval. Weight values are relative to the initial value before learning.

Next, we investigated the activation of assemblies that had undergone intermediate consolidation after being learned, meaning that the breaks shown in Fig. 1d lasted for 8 hours instead of 3 seconds. Although LTD was switched on for these investigations, much of the learning order effect vanishes as well (Fig. 4b,d). The effect of hub-like structures is also similar to the no-LTD case. It seems that here, LTD is not significantly expressed in the synapses within the assemblies, even though LTD occurs in other synapses (cf. Fig.4b and inset in Fig. 4a). Apparently, if only one assembly is learned before the next one follows after hours, much of the depression which evokes the learning order effect is too weak to become consolidated (also cf. Suppl. Figs. S1 and S4).

These results indicate that diverse effects arise from the interaction of plasticity processes on different timescales, triggered by different learning/consolidation protocols. While the standard protocol causes both late-phase LTP and substantial late-phase LTD, leading to the described strong learning order effect and its reduction for assemblies with less overlaps, the intermediate consolidation protocol rather causes operation in the regime of late-phase LTP, leading to a weaker learning order effect and promoted activation of hub assemblies.

### Priming on long timescales

After investigating the functional implications of different organizational paradigms, in this subsection, we provide a proof of principle to show that our two-phase plasticity model is able to describe priming on long timescales. To this end, we use a protocol that includes a brief additional stimulus delivered to all neurons of one selected assembly after learning and consolidation (Fig. 5a). This stimulation causes the respective assembly to be transiently primed for later reactivation, which we investigated by considering the likelihood of avalanches under background noise, as it was done in the previous section. Our results demonstrate a mechanism for priming enabled by the interplay of early- and late-phase long-term plasticity (Fig. 5b-e). The priming decays on a timescale of minutes to hours. Interestingly, in the non-overlapping paradigm, the priming stimulation causes the primed memory representation to behave in an almost attractor-like manner (Fig. 5b, where the likelihood of activation reaches values of more than 0.4).

**Figure 5:**
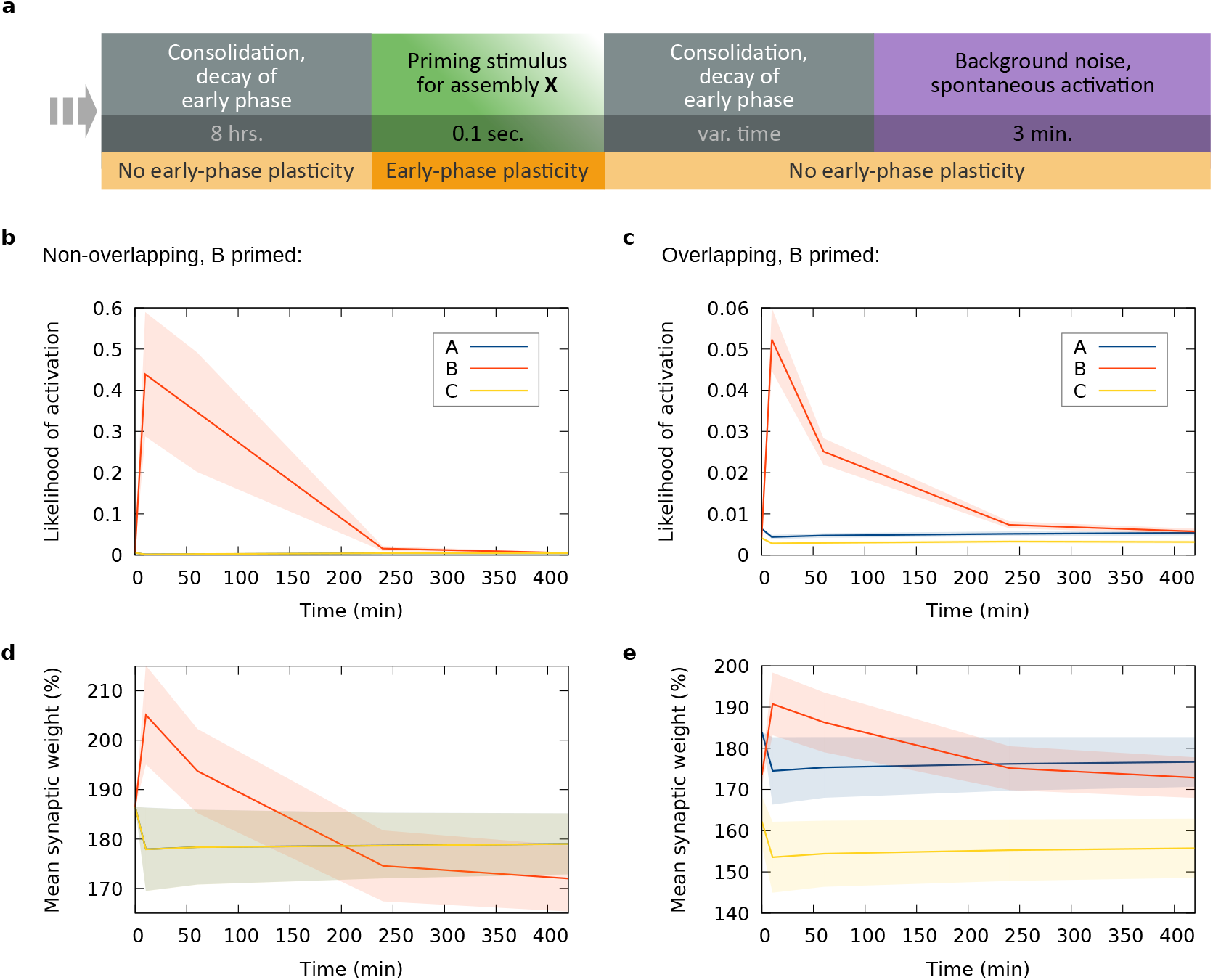
Priming on long timescales is enabled by synaptic tagging and capture. **(a)** Consolidation and priming phase (following learning as sketched in Fig. 1d, with order *A*-*B*-*C*-*A*-*C*-*B*-*B*-*A*-*C*-*A*-*B*-*C*-*C*-*B*-*A*): consolidation takes place for eight hours after learning. After that, a priming stimulus is applied to one of the assemblies, followed by another period of consolidation and early-phase decay. Then, driven by background noise, spontaneous activity of the memory representations is observed for three minutes. **(b,c)** Resulting likelihood of avalanches in the three assemblies at different times after priming *B*. The values at time zero show the case before (without) priming. **(d,e)** Temporal development of the mean synaptic weight in the assemblies, for the same cases as in **b** and **c**. Data in panels **b-e** are averaged over 10 networks. Error bands show standard error of the mean. Weight values are relative to the initial value before learning.

For investigating priming in the absence of other effects, we sought to attenuate the learning order effect and therefore used the extended interleaved protocol introduced in the previous section. For the non-overlapping case, this approach works fine as it almost entirely eliminates the influence of the learning order (Fig. 5b). However, in the overlapping case (Fig. 5c), the learning order effect still plays a role, since in the basal state, *A* is much more active than *B* and *C*. The priming of *B* shows a significant effect in the first hour, but then the effect is overshadowed by the high activity of assembly *A*. Finally, we have to mention that in rare cases, priming stimulation after learning with the interleaved protocol seems to disrupt the recallable memory structure and render reactivation impossible (this occurred in 1 of 10 trials).

The mean weight plotted in Fig. 5d,e shows roughly the same qualitative time course as the activation (Fig. 5b,c). It should be noted that in Fig. 5d, the mean weight in assembly *B* decreases to a lower value than it had before priming, which can be explained by the strong depression of a few synapses that thus enter the late phase, while most synapses are subject to early-phase potentiation.

Data for more paradigms are shown in Suppl. Fig. S7. The case where only 50% of the assembly neurons receive the priming stimulus is shown in Suppl. Fig. S8 (more paradigms in Suppl. Fig. S9). This case leads to the counterintuitive but interesting outcome of priming by differential degree of LTD. Further investigations on this are, however, beyond the scope of this study.

## Discussion

In this study, we have shown by proof of principle that the molecular mechanisms of calcium-based synaptic plasticity and STC enable to robustly learn, consolidate, organize, and prime long-term memory representations. Our results demonstrate that these features give rise to specific characteristics as they are found in neuropsychological experiments.

Measuring the likelihood of spontaneous activation of three cell assemblies in a spiking recurrent neural network after learning and consolidation, we found two factors that guide the activation: 1. the learning order, and 2. the type of overlaps between the assemblies. As an experimental study has shown, overlap accounts for memory association but is not necessary for the recall of the original separate memories, which matches our findings for the non-overlap and overlap paradigms [54]. Moreover, there is experimental evidence that memories can be potentiated and depotentiated independently of each other [55], which also seems to be possible in our model. Data from free recall experiments typically exhibit primacy (high probability of activating the first learned items), recency (high probability of activating the last learned items), and contiguity (higher probability of transition between neighboring items) [31, 33]. By our model, we provide a mechanistic explanation for the recency effect. The direct reproduction of data from free recall experiments [34, 56, 57] with our model would, however, require a larger number of cell assemblies, which was beyond the capacity of this study. Since our detailed network model had a high demand of computational power, we had to restrict our investigations to a medium-sized network of 3125 neurons and three assemblies. Nevertheless, direct matching of a larger network based on our model and data from free recall experiments could be the goal of a future study. Extending our model to more assemblies would also be very helpful for investigating and modeling contiguity, complementary to previous models [28, 29]. Another interesting approach in a future study could be to adjust our network such that the memory representations constitute attractors. This would enable the study of the transitions between attractor states, which have been proposed to account for the switching between concepts [29, 58, 59], and would build on the vast amount of previous studies on attractor neural networks.

In addition to recency in free recall, our model may provide a mechanistic basis for retroactive interference [48, 49, 50] and enhanced recall of similar memories [51, 52]. Our findings show that the activation of memory representations featuring hub-like overlaps is enhanced or attenuated, depending on the consolidation protocol (Figs. 3c and 4d). Moreover, for longer intervals between learning stimuli but otherwise equal parameters, “memory-improving” interference occurs (Suppl. Fig. S6) instead of the “memory-deteriorating” interference that results from the standard protocol (Suppl. Fig. S5). From our theoretical investigations, we made the prediction that the effects described above are essentially caused by LTD (Fig. 4a,c). This can be tested experimentally by a protocol that blocks LTD, for example, via the mGlu receptor antagonist MCPG, or via BDNF [60, 61]. How our findings relate to experiments showing that the acquisition of overlapping assemblies sufficiently close in time facilitates recall, while memories acquired further apart in time tend to be separated [62, 63], remains to be shown in future investigations. Furthermore, the experimental investigation of our “intermediate consolidation” protocol could yield a large gain in knowledge. The difference between this protocol and typical free recall protocols is that after learning a memory representation we wait until it is consolidated (i.e., for about 8 hours), and then learn the next representation, whereas in free recall experiments, the whole list of items is usually learned in short time before consolidation may take place (similar to our “standard” protocol). The experimental investigation of the “intermediate consolidation” protocol would enable to test our predictions and yield further interesting insights in the interaction of short and long-term memory for free recall. An issue could be that the subjects would have to stay awake for the whole duration of the protocol of about 24 hours, to avoid a biasing effect from sleep consolidation. However, similar experiments have already been performed to test the impact of sleep deprivation on free recall [39, 64].

In our simulations, we assumed early-phase plasticity to be disabled during certain phases. As we pointed out in the Methods section, this is justified by a lack of novelty or attention, which goes along with lowered neuromodulator concentrations [65, 66], and thereby prevents learning [67, 68]. If, however, early-phase plasticity were enabled all the time, spontaneous activity could exert the following influence on the structure of a cell assembly: either the assembly would stay unaffected, as we have shown for smaller cell assemblies in our previous study [9], or, LTD would cause a degradation by downregulating the synaptic weights, or, LTP would cause reinforcement by increasing the synaptic weights. While the latter is not possible due to the firing rate threshold in our model, we assume that spontaneous occurrence of LTD, if any, would not qualitatively change our results.

In the second part of our study, we demonstrated the priming of one of the three previously learned and consolidated memory representations, on a timescale of minutes to hours. This effect is essentially caused by the interplay between early- and late-phase long-term plasticity. After consolidation, the structure of the memory representations is permanently stored by the late-phase weights, while additional transient information, like priming information, is stored by the early-phase weights, which decay across minutes to hours. However, as our results also indicate, priming information may “spill over” to the late phase, causing permanent changes in the structure of a memory representation. In our study, the priming effect is strongest if priming is only 10 minutes ago, and declines until it has vanished almost completely after 7 hours (Fig. 5b,c). This time dependence, as well as the impact of priming in general, may be affected by the strength and duration of the priming stimulus which “loads the representation into short-term memory”. Moreover, the effect of the time delay may be influenced by a wide range of further experimental conditions, making it difficult to draw a general conclusion, which was pointed out by others earlier [69]. As our results constitute a proof of principle, we leave it to further studies to investigate the influence of different kinds of stimulation. Furthermore, while we showed that our model can account for direct positive priming, our model can likely be extended with relatively low effort to account for other types of priming. We already encountered predominance of LTD in the case where only 50% of the assembly neurons received priming stimulation (Suppl. Figs. S8 and S9). While this paradigm still causes a slightly positive priming effect, lowering this fraction even further or simply applying background activity in the presence of early-phase plasticity should revert the outcome and cause a negative priming effect. Such further results would be of great importance, especially because a mechanistic biological theory for negative priming still remains to be found [44, 70, 71]. Moreover, a tuned version of our model could possibly account for semantic priming, which would be the case if the stimulation of an assembly that overlaps with a second assembly would cause primed activation of the second assembly (e.g., in the way that the word “chicken” would prime the word “duck”) [45, 69, 72, 73]. Finally, the fact that the interplay of early- and late-phase plasticity enables priming is another hint that they might constitute correlates for the concepts of retrieval strength and storage strength [9, 74].

To summarize, we could show that LTD on the one hand and the interplay of early and late-phase long-term plasticity on the other hand exert crucial influence on memory function. Our results provide neural correlates to – at least partially – explain different behavioral effects. In the future, our model may serve as a mechanistic basis for the detailed reproduction of findings obtained in behavioral experiments.

## Methods

### Model

To simulate the dynamics of memory representations, we used a network model that comprises spiking neurons and synapses with detailed plasticity features. In this section, we first present our mathematical description of neurons and synapses, which is depicted in Fig. 1a, as well as the structure of our network at the population level. After that, we explain the different overlap paradigms and stimulation protocols.

The parameters that we used are given in Tables 1 and 2. Parts of this section have been adapted from our previous study [9], which provided the basis for our model.

**Table 1:** Parameters for neuron and static network dynamics. Values were used as given in this table, unless stated otherwise.

| Symbol | Value | Description | Refs. |
| --- | --- | --- | --- |
| $\Delta t$ | 0.2 ms | Duration of one time step for numerical computation | this study |
| $\tau_m$ | 10 ms | Membrane time constant | [75, 76] |
| $\tau_{\text{syn}}$ | 5 ms | Synaptic time constant, also for external input current | [75, 76, 77] |
| $t_{\text{ax,delay}}$ | 3 ms | Axonal spike delay | [76, 78] |
| $t_{\text{ref}}$ | 2 ms | Duration of the refractory period | [75, 79] |
| $R_m$ | 10 MΩ | Membrane resistance | [75] |
| $V_{\text{rev}}$ | −65 mV | Reversal (equilibrium) potential | [75] |
| $V_{\text{reset}}$ | −70 mV | Reset potential | [75] |
| $V_{\text{th}}$ | −55 mV | Threshold potential to be crossed for spiking | [75] |
| $I_0$ | 0.15 nA | Mean of the external background current | [9] |
| $\sigma_{\text{wn}}$ | 0.05 nA s<sup>1/2</sup> | Standard deviation for Gaussian noise in the background current | [9] |
| $N_e$ | 2500 | Number of neurons in the excitatory population | this study |
| $N_i$ | 625 | Number of neurons in the inhibitory population | this study |
| $p_c$ | 0.1 | Probability of a connection existing between two neurons | [11] |
| $h_0$ | 4.20075 mV | Initial excitatory→excitatory coupling strength | [9, 80] |
| $w_{ei}$ | 2 $h_0$ | Excitatory→inhibitory coupling strength | [9] |
| $w_{ie}$ | 3.5 $h_0$ | Excitatory→inhibitory coupling strength | this study |
| $w_{ii}$ | 3.5 $h_0$ | Inhibitory→inhibitory coupling strength | this study |
| $f_{\text{stim}}$ | 100 Hz | Frequency of stimulation | [80, 81] |
| $N_{\text{stim}}$ | 25 | Number of input neurons for stimulation | [9] |
| $n_{\text{thresh}}$ | 10 | Detection threshold for an avalanche | this study |

**Table 2:**
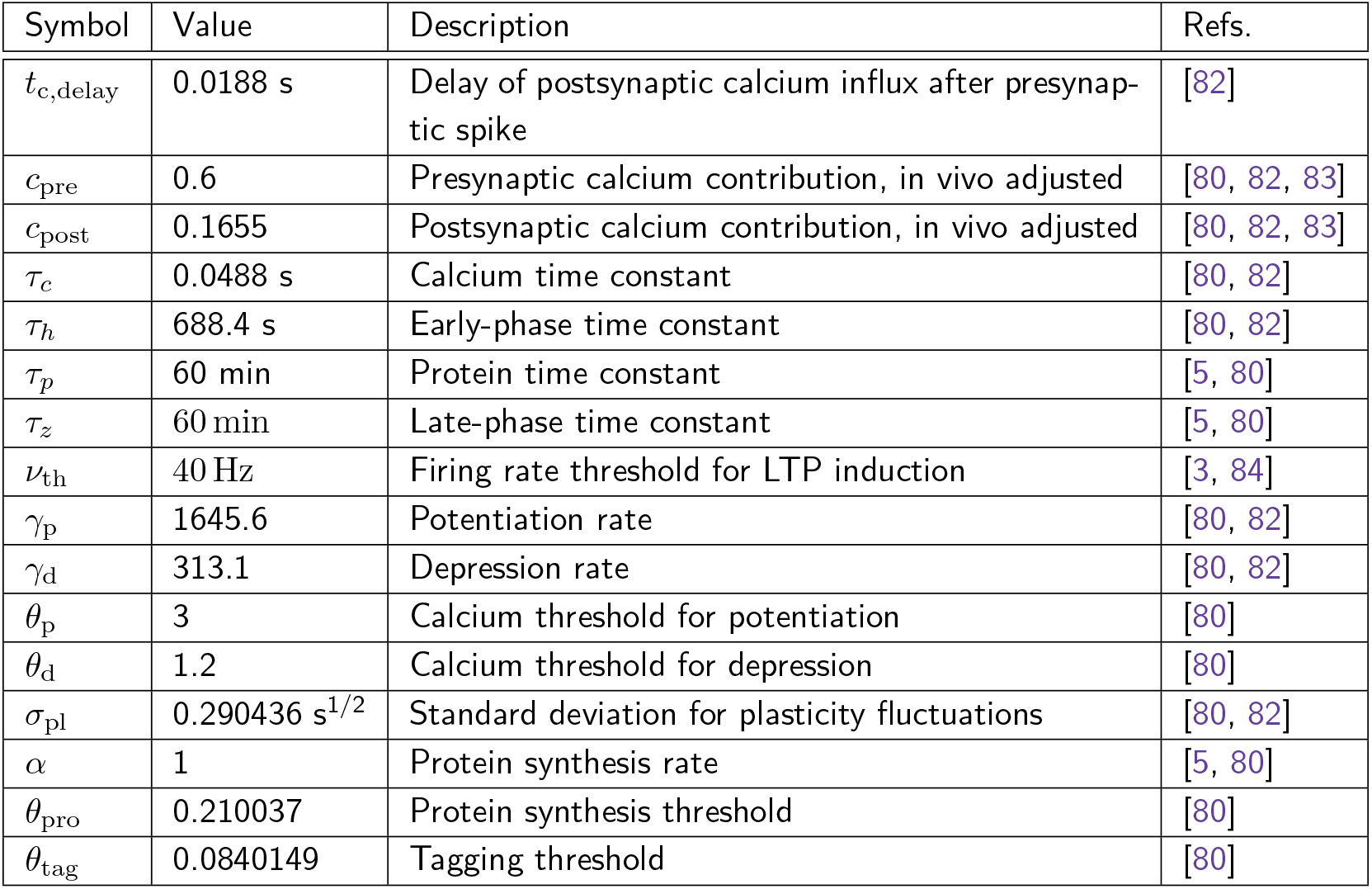
Parameters for synaptic plasticity. Values were used as given in this table, unless stated otherwise.

### Neuron model

The dynamics of the membrane potential *V_i_*(*t*) of the Leaky Integrate-and-Fire neuron *i* is described by [76]:

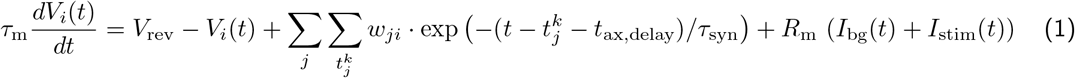

with reversal potential *V*_rev_, membrane time constant *τ*_m_, membrane resistance *R*_m_, synaptic weights *w_ji_*, spike times 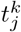, axonal delay time *t*_ax,delay_, synaptic time constant *τ*_syn_, external background current *I*_bg_(*t*), and external stimulus current *I*_stim_(*t*). If *V_i_* crosses the threshold *V*_th_, a spike is generated. The spike time 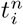 is then stored and the membrane potential is reset to *V*_reset_, where it remains for the refractory period *t*_ref_. Under basal conditions, the membrane potential dynamics is mainly driven by an external background current that accounts for synaptic inputs from outside the network, described by an Ornstein-Uhlenbeck process:

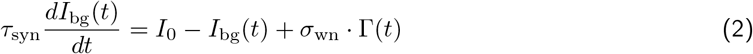

with mean current *I*_0_ and white-noise standard deviation *σ*_wn_. In this equation, Γ(*t*) is Gaussian white noise with mean zero and variance 1/*dt* that approaches infinity for *dt* → 0. The Ornstein-Uhlenbeck process has the same colored-noise power spectrum as the fluctuating input to cortical neurons coming from a large presynaptic population, and is therefore well-suited to model the background noise in our model [21].

In addition to the background noise, we use another Ornstein-Uhlenbeck process to model the stimulus current *I*_stim_(*t*) for learning, recall, and priming (cf. the subsection ‘Learning, recall, and priming stimulation’ below):

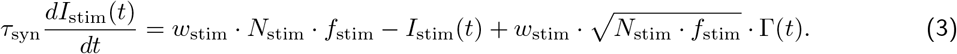

Mean and standard deviation of this process are defined by putative input spikes from *N*_stim_ neurons, occurring at the frequency *f*_stim_ and conveyed by synapses of weight *w*_stim_ to the neurons of our network that are to be stimulated.

### Model of synapses and STC

All spikes *k* that occur in neuron *j* are transmitted to *i*, if there is a synaptic connection from neuron *j* to neuron *i*. The postsynaptic current caused by a presynaptic spike then depends on the total weight or strength of the synapse, which is is given by:

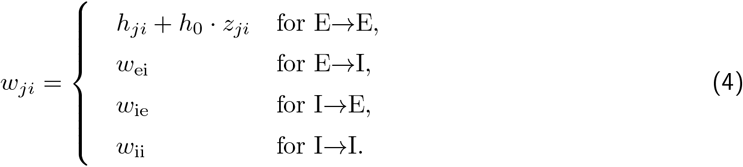

While in the model, all synaptic connections involving inhibitory neurons are constant, the weight of E→E connections comprises two variable contributions to account for STC: the early-phase weight *h_ji_*, and the late-phase weight *z_ji_*. The dynamics of the early-phase weight is given by

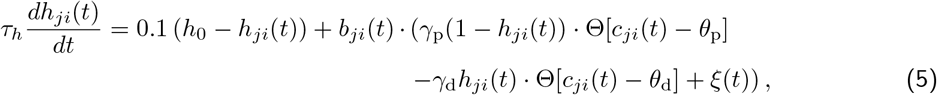

where Θ[*·*] is the Heaviside function, *τ_h_* is a time constant, *c_ji_*(*t*) is the calcium concentration at the postsynaptic site, and *b_ji_*(*t*) describes special additional conditions for the induction of plasticity. The first term on the right-hand side describes the decay of the early-phase weight to its initial value *h*_0_, the second term describes early-phase LTP with rate *γ*_p_, given that calcium is above the threshold *θ*_p_, the third term describes early-phase LTD with rate *γ*_d_, given that calcium is above the threshold *θ*_d_, and the fourth term 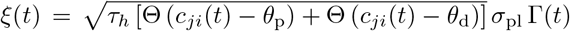 describes calcium-dependent fluctuations with standard deviation *σ*_pl_ and Gaussian white noise Γ(*t*) with mean zero and variance 1/*dt*. The calcium concentration *c_ji_*(*t*) at the postsynaptic site is driven by the pre- and postsynaptic spikes *n* and *m*, respectively:

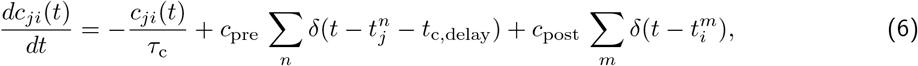

where *δ*(·) is the Dirac delta distribution, *τ_c_* is a time constant, *c*_pre_ is the contribution of presynaptic spikes, *c*_post_ is the contribution of postsynaptic spikes, 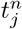 and 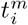 are spike times, and *t*_c,delay_ is the delay *j*_*i*_ of calcium triggered by presynaptic spikes.

In addition to the calcium dependence of early-phase plasticity, which is based on previous studies [9, 80, 82, 83]), we imposed special conditions for the occurrence of plasticity to enable the stable formation, consolidation, and activation of large cell assemblies. In their original study [82], Graupner and Brunel had only considered firing rates up to 50 Hz. A drawback in their calcium-based model is the chance that spiking at high rates on only one side of the synapse would suffice to trigger LTP, which is physiologically not plausible. To account for the evidence that the firing of only one neuron (i.e., on one side of the synapse) at high rates should not be sufficient to evoke LTP [12, 14, 85], we introduced an additional threshold *ν*_th_ which allows the induction of LTP only if both pre- and postsynaptic firing rate cross this threshold. Furthermore, we blocked early-phase plasticity outside the learning and priming phases to enable fast computation and stable activation dynamics. This is explained by a lack of novelty outside the learning and priming phases, which goes along with lowered neuromodulator concentrations [65, 66] and thereby prevents learning, as experimental studies have demonstrated [67, 68]. In summary, the two “special plasticity conditions” that we added to the model can be put down formally in the following way:

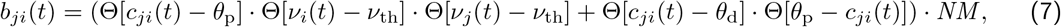

where the first term in the round brackets describes the LTP induction depending on the calcium concentration *c_ji_* and the firing rate threshold *ν*_th_, and the second term describes the LTD induction which only depends on calcium. The firing rate function *ν_i_* (*t*) describes the firing rate of neuron *i* at time *t*, computed using a sliding window of 0.5s. The factor *NM* is the neuromodulatory impact, which we assume unity during learning and priming phases and zero otherwise.

Driven by the calcium-based early-phase dynamics, the late-phase synaptic weight is given by:

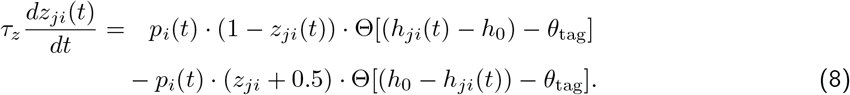

with the protein amount *p_i_*(*t*), the late-phase time constant *τ_z_*, the maximum late-phase weight *z*_max_, and the tagging threshold *θ*_tag_. The first term on the right-hand side describes late-phase LTP, while the second one describes late-phase LTD. Both depend on the amount of proteins being available. If the early-phase weight change *|h_ji_*(*t*) *− h*_0_| exceeds the tagging threshold, the synapse is tagged. This can be the case either for positive or for negative weight changes. The presence of the tag leads to the capture of proteins (if *p_i_*(*t*) > 0), and thereby gives rise to changes in the late-phase weight.

The synthesis of new proteins requires sufficient changes in early-phase weights, while there is also an inherent decay of the protein amount [5]:

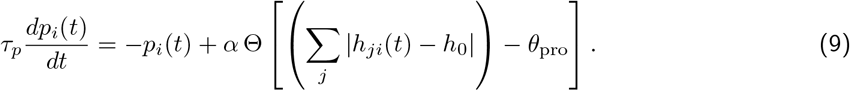

### Population structure

Using the neuron model and the synapse model explained above, we set up a neural network consisting of 2500 excitatory and 625 inhibitory neurons (depicted in Fig. 1b). The ratio of 4:1 between excitatory and inhibitory neurons is typical for cortical and hippocampal networks [86]. Some of the excitatory neurons receive specific inputs for the formation of cell assemblies (see the subsection ‘Learning, recall, and priming stimulation’). The inhibitory population provides feedback inhibition, while not directly being subject to plastic changes. The probability for the existence of a synapse between two neurons is 0.1, across the whole network. This is a plausible value for hippocampal region CA3 [11]. For every trial, a new network structure was drawn.

### Learning, recall, and priming stimulation

Before we apply the first learning stimulus to our network, we let the initial activity settle for 10.0 seconds. Then, we apply our learning protocol, which delivers three stimulus pulses, of 0.1 seconds duration each, to the neurons dedicated to the first cell assembly (*A*, the first 600 neurons in the network). Breaks of 0.4 seconds separate the pulses. The stimulation is delivered by an Ornstein-Uhlenbeck process (see Eq. 3 introduced above). After the last pulse has ended, we let the network run under basal conditions for 3.0 seconds (Fig. 1d). For the case of intermediate consolidation, which we consider in one part of our study, we extend this period to 28800.0 seconds, such that the synaptic weights may undergo consolidation through synaptic tagging and capture. Every time that after stimulation ends, We disable early-phase plasticity (except for the decay of early-phase changes) after a time window of 12.5 seconds. This is reasonable considering the impact of novelty and neuromodulation on encoding [67, 68].

Then, we apply another learning stimulus of the same kind as the first one to the neurons dedicated to the second cell assembly *B*. In the non-overlapping case, this set of neurons is completely disjunct from the first set. In the cases with overlaps, 10% of the neurons in the assembly (in the main paper; also cf. Suppl. Table S1) are shared with the other assembly. The size of the overlaps is of the same order of magnitude as experimental findings, which range from around 1% to more than 40% [54, 62, 87, 88]. After the learning stimulus, we let the network again run under basal conditions for 3.0 seconds (28800.0 seconds in the intermediate consolidation case).

Finally, we apply a third learning stimulus of the same kind, this time to the neurons dedicated to the third cell assembly *C*. In the non-overlapping case, this set of neurons is completely disjunct from the first and second set. For the overlap cases, still, 10% of the neurons between each two assemblies are shared. We denote the exclusive intersections between each two assemblies by *I_AB_*, *I_AC_*, *I_BC_* (see Fig. 1c), which make up for half of each overlap between two assemblies. The intersection between all three assemblies *I_ABC_* makes up for the other half of each overlap between two assemblies (cf. Suppl. Table S1). Specifically, the different subpopulations are set to occupy the following neuron indices in the network (example for overlapping case/“OVERLAP10”):

- *I_ABC_*: 540–569,
- *I_AC_*: 0–29,
- 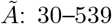,
- *I_AB_*: 570–599,
- 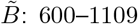,
- *I_BC_*: 1110–1139,
- 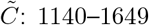.

By *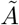*, *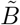*, and *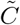* we denote the subsets of neurons that exclusively belong to one of the assemblies. Note that in the non-overlapping case, 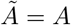, 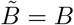, and 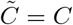.

At two points in our study, we used interleaved learning of the three memory representations. For this, instead of the three steps A-B-C, we employed the following scheme of 15 learning steps: *A*-*B*-*C*-*A*-*C*-*B*- *B*-*A*-*C*-*A*-*B*-*C*-*C*-*B*-*A*.

After learning, we let the network first run under basal conditions for 3.0 seconds and then consolidate for 28800.0 seconds (eight hours, see Fig. 1e). This caused all early-phase weight changes to either decay or be transferred to the late phase, thereby yielding long-term memory representations. After that, we started measuring the spontaneous activity (see section below).

For the testing of memory function in Fig. 2, we applied a recall stimulus to randomly drawn 20% of the neurons of an assembly, delivered by one pulse 0.1 seconds long, with otherwise the same parameters as for the learning stimulus. During this, we left early-plasticity switched off.

For the priming of a memory representation, which we did in the last part of our study, we applied a stimulus to all neurons of one selected assembly (alternatively, to randomly drawn 50% of the assembly neurons, the results of which are only shown in the Supplementary Information). The stimulus was delivered via one pulse 0.1 seconds long, with otherwise the same parameters as for the learning stimulus (Fig. 5a). The priming stimulus was followed by a period of consolidation and early-phase decay of varied duration, before we started measuring the spontaneous activity.

### Determining avalanches in the spontaneous activity

After the learning, consolidation, and possibly priming of the cell assemblies, we measured the network activity across three minutes to evaluate the spontaneous activity of the assemblies. To this end, we computed the likelihood that an avalanche with a certain minimum size occurred in a specific assembly, averaged across the whole period of three minutes. In addition, we measured the plain mean firing rate of each assembly and the control population across the whole period of three minutes.

To determine if an avalanche occurs, we counted the number of spikes that were produced by the neurons of each cell assembly in a time frame of 10 ms. Then, we compared these numbers to the threshold value *n*_thresh_, and considered an assembly active if the threshold was reached. Multiple assemblies could be active at the same time. Then, we computed the likelihood of an avalanche in assembly *A*, for instance, by dividing the number of periods *N*_act_(*A*) in which the assembly was active by the total number of time frames (given by the total duration of the simulation *t*_max_/10 ms):

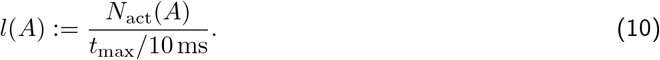

The likelihoods for avalanches in *B* and *C* are computed analogously. The threshold value *n*_thresh_ was chosen to be about the border value of the 99% quantile for the number of spikes of an assembly in one time frame (Suppl. Fig. S10).

### Computational implementation and software used

We used C++ in the ISO 2011 standard to implement our simulations. To compile and link the code, we employed g++ in version 7.5.0 with boost in version 1.65.1.

For further data analysis we used Python 3.7.3 with NumPy 1.20.1 and pandas 1.0.3. For the creation of plots we used gnuplot 5.0.3, as well as Matplotlib 3.1.0.

We generated random numbers using the generator ‘minstd_rand0’ from the C++ standard library, while the system time served as the seed. We implemented a loop in our code which ensured that for each distribution a unique seed was used. The simulations that had to be performed for this study were computationally extremely demanding. Thus, it was very fortunate that we had the opportunity to run most of our jobs on the computing cluster of the Gesellschaft für wissenschaftliche Datenverarbeitung mbH Göttingen (GWDG).

## Data availability

All data underlying the results presented in this study are contained in this manuscript and the Supporting Information. The data can be reproduced using the simulation code and the analysis scripts that we have released [89].

## Code availability

Our code is freely available and has been released under the Apache-2.0 license [89]. As it continues to be developed, the latest version can be retrieved from https://github.com/jlubo/memory-consolidation-stc.

## Acknowledgments

We thank the members of the Department of Computational Neuroscience, especially Sebastian Schmitt, for many helpful comments on this study. The research was funded by the German Research Foundation (CRC1286, project C1, project #419866478) and by the H2020 - FETPROACT project Plan4Act (#732266).

## Author contributions

Conceptualization, methodology, and investigation: CT and JL; software and simulations: JL; data curation, formal analysis, and visualization: JL; funding acquisition and supervision: CT; writing and editing: CT and JL.

