## Supplementary Information for "Organization and priming of long-term memory representations with two-phase plasticity"

2021-04-15

[1] Department of Computational Neuroscience, III. Institute of Physics – Biophysics, University of Göttingen, Göttingen, Germany

[2] Bernstein Center for Computational Neuroscience, Göttingen, Germany

Contact:
[*]
[#]

| Label | Appears in main paper | Description |
|---|---|---|
| NOOVERLAP | x | No overlaps |
| OVERLAP10 | x | Each two assemblies exclusively overlap by 5%, and all three assemblies commonly overlap by 5% |
| OVERLAP10 no ABC | | Each two assemblies exclusively overlap by 10%, there is no common overlap between all three assemblies |
| OVERLAP10 no AC, no ABC | x | Both $A$, $B$ and $B$, $C$ overlap by 10% |
| OVERLAP10 no BC, no ABC | x | Both $A$, $B$ and $A$, $C$ overlap by 10% |
| OVERLAP15 | | Each two assemblies exclusively overlap by 7.5%, and all three assemblies commonly overlap by 7.5% |
| OVERLAP15 no ABC | | Each two assemblies exclusively overlap by 15%, there is no common overlap between all three assemblies |
| OVERLAP15 no AC, no ABC | | Both $A$, $B$ and $B$, $C$ overlap by 15% |
| OVERLAP15 no BC, no ABC | | Both $A$, $B$ and $A$, $C$ overlap by 15% |
| OVERLAP20 | | Each two assemblies exclusively overlap by 10%, and all three assemblies commonly overlap by 10% |
| OVERLAP20 no ABC | | Each two assemblies exclusively overlap by 20%, there is no common overlap between all three assemblies |
| OVERLAP20 no AC, no ABC | | Both $A$, $B$ and $B$, $C$ overlap by 20% |
| OVERLAP20 no BC, no ABC | | Both $A$, $B$ and $A$, $C$ overlap by 20% |

Supplementary Table S1: All organizational paradigms referred to by the Supplementary Information.

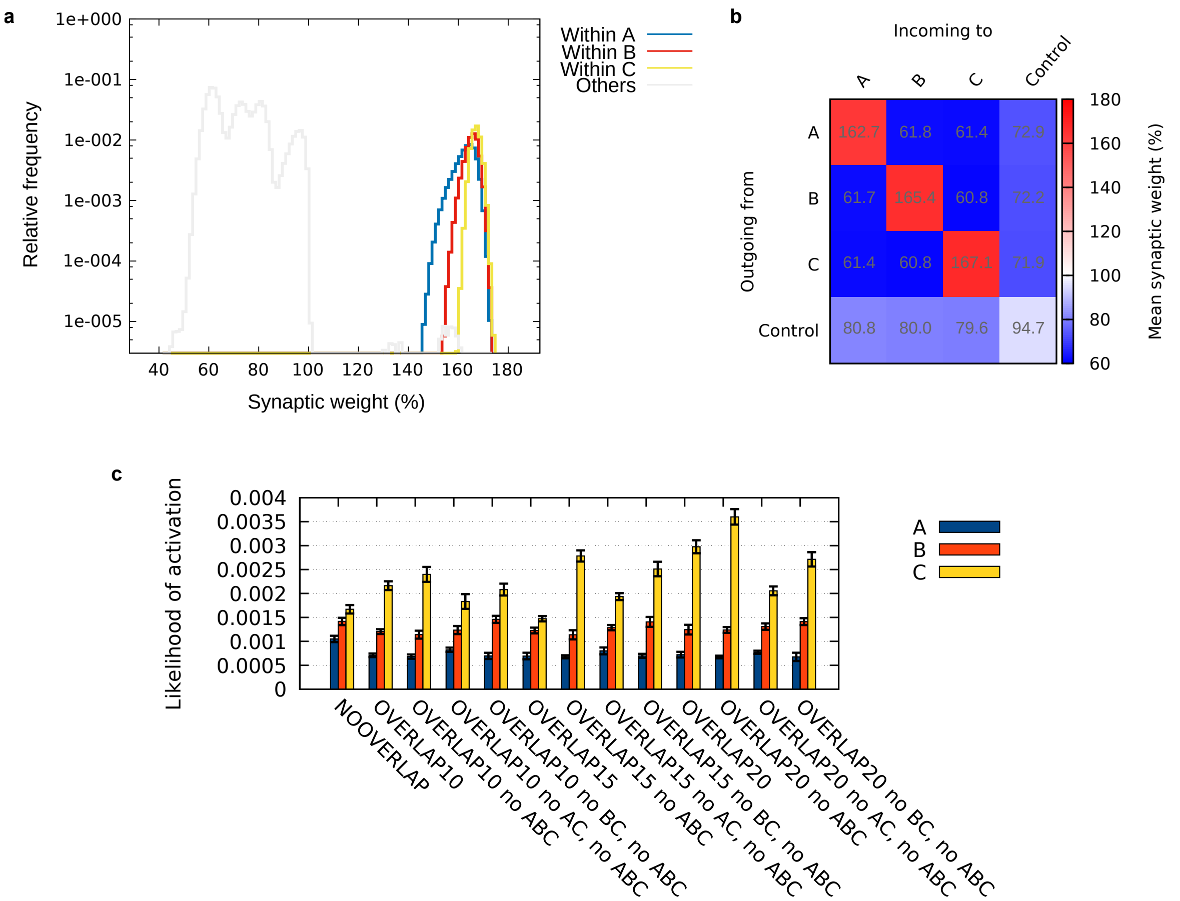


Supplementary Figure S1: NOOVERLAP paradigm; after learning but before consolidation (standard protocol, learning order: $A$-$B$-$C$). **(a)** Weight distribution revealing the spread of LTP and LTD. **(b)** Abstract weight matrix showing the mean weight within and between all subpopulations of the excitatory population: the exclusive parts of the cell assemblies, the exclusive intersections, and the control population. The asymmetric nature of the matrix reveals the directionality of the coupling strengths. **(c)** During spontaneous activity, driven by background noise: overview of likelihood of avalanches in the memory representations across different organizational paradigms, including overlaps of varying size. Error bars show the 95% confidence interval. Data in all panels were averaged over $10$ trials. Weight values are relative to the initial value before learning.

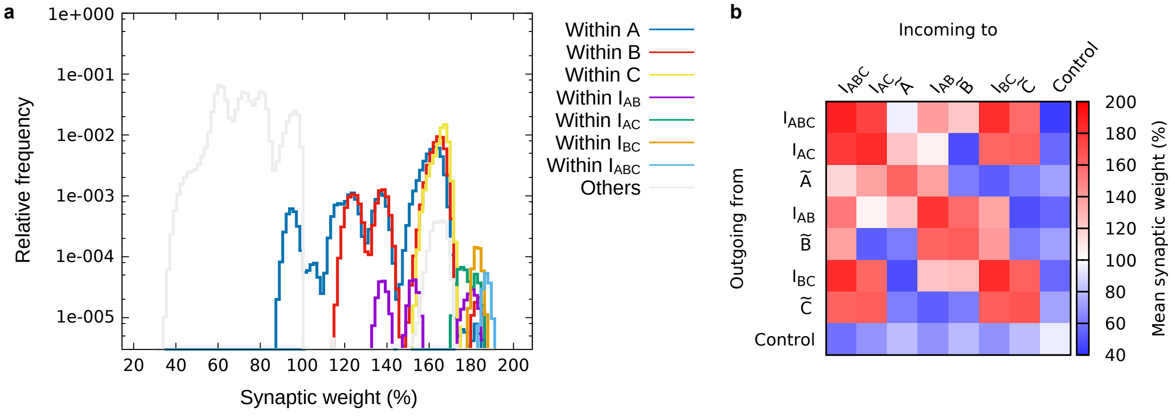


Supplementary Figure S2: OVERLAP10 paradigm; after learning but before consolidation (standard protocol, learning order: $A$-$B$-$C$). **(a)** Weight distribution revealing the spread of LTP and LTD. **(b)** Abstract weight matrix showing the mean weight within and between all subpopulations of the excitatory population: the exclusive parts of the cell assemblies, the exclusive intersections, and the control population. The asymmetric nature of the matrix reveals the directionality of the coupling strengths. Data in both panels were averaged over $10$ trials. Values are relative to the initial value before learning.

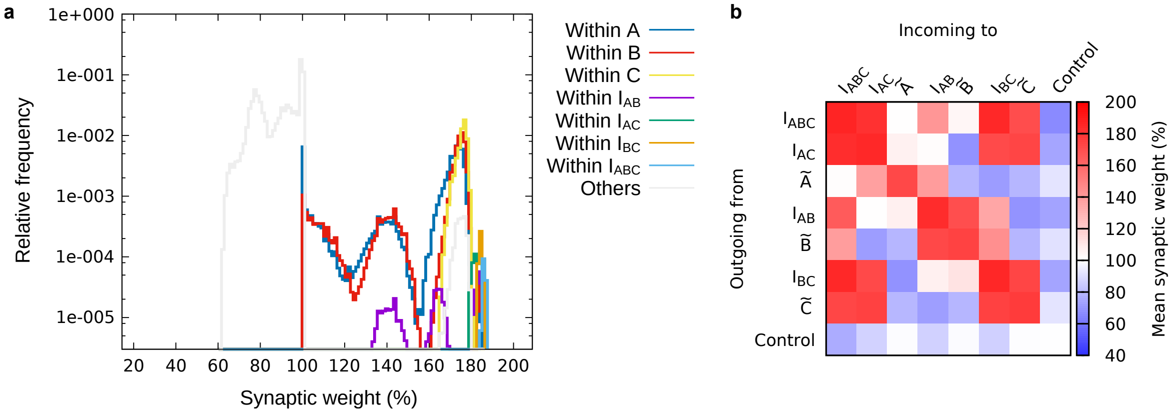


Supplementary Figure S3: OVERLAP10 paradigm; after learning and consolidation (standard protocol, learning order: $A$-$B$-$C$). **(a)** Weight distribution revealing the spread of LTP and LTD. **(b)** Abstract weight matrix showing the mean weight within and between all subpopulations of the excitatory population: the exclusive parts of the cell assemblies, the exclusive intersections, and the control population. The asymmetric nature of the matrix reveals the directionality of the coupling strengths. Data in both panels were averaged over $10$ trials. Values are relative to the initial value before learning.

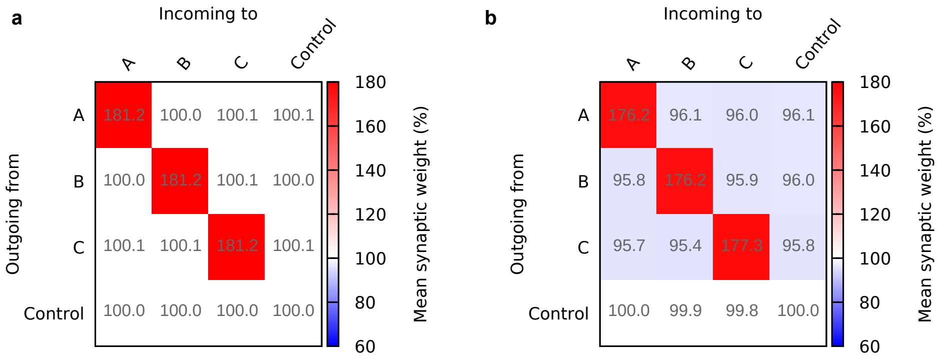


Supplementary Figure S4: Abstract weight matrices showing the mean weight within and between all subpopulations of the excitatory population after learning and consolidation (order: $A$-$B$-$C$). **(a)** NOOVERLAP paradigm with blocked LTD, **(b)** NOOVERLAP paradigm with intermediate consolidation of $8$ hours in between learning $A$, $B$, and $C$. Data in both panels were averaged over $10$ trials. Values are relative to the initial value before learning.

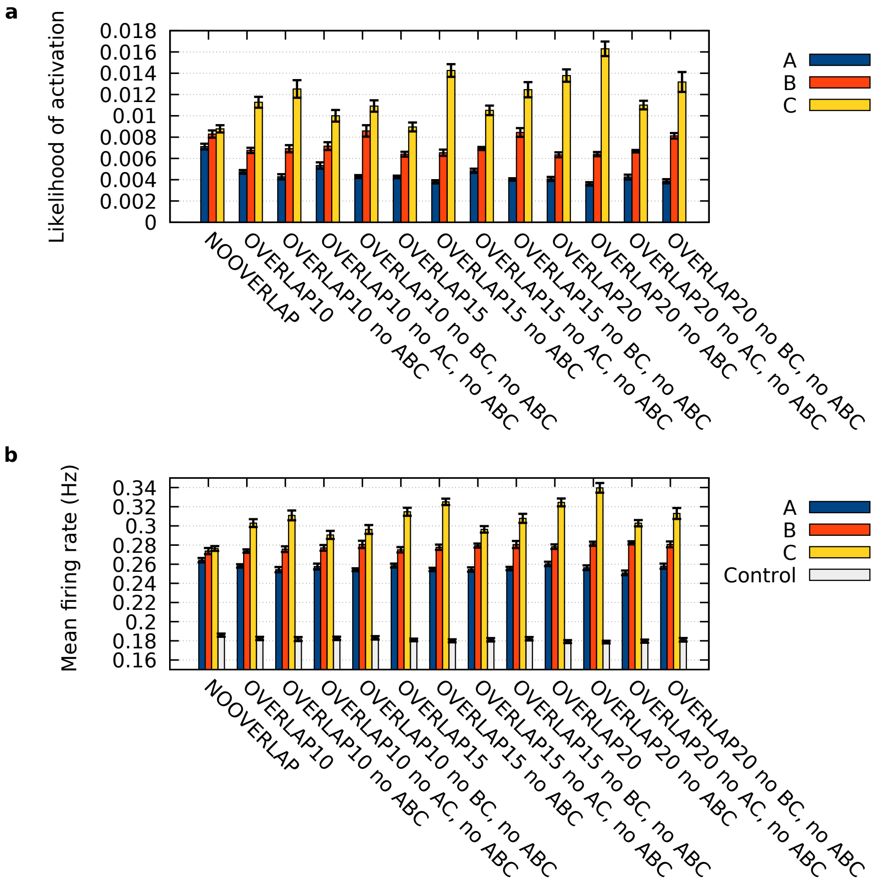


Supplementary Figure S5: Spontaneous activity of long-term memory representations, driven by background noise. Standard protocol for learning/consolidation (order: $A$-$B$-$C$). **(a)** Overview of likelihood of avalanches across different organizational paradigms, including overlaps of varying size. **(b)** Overview of mean firing rates in the same organizational paradigms as in **a**. Averaged over $10$ trials. Error bars show the 95% confidence interval.

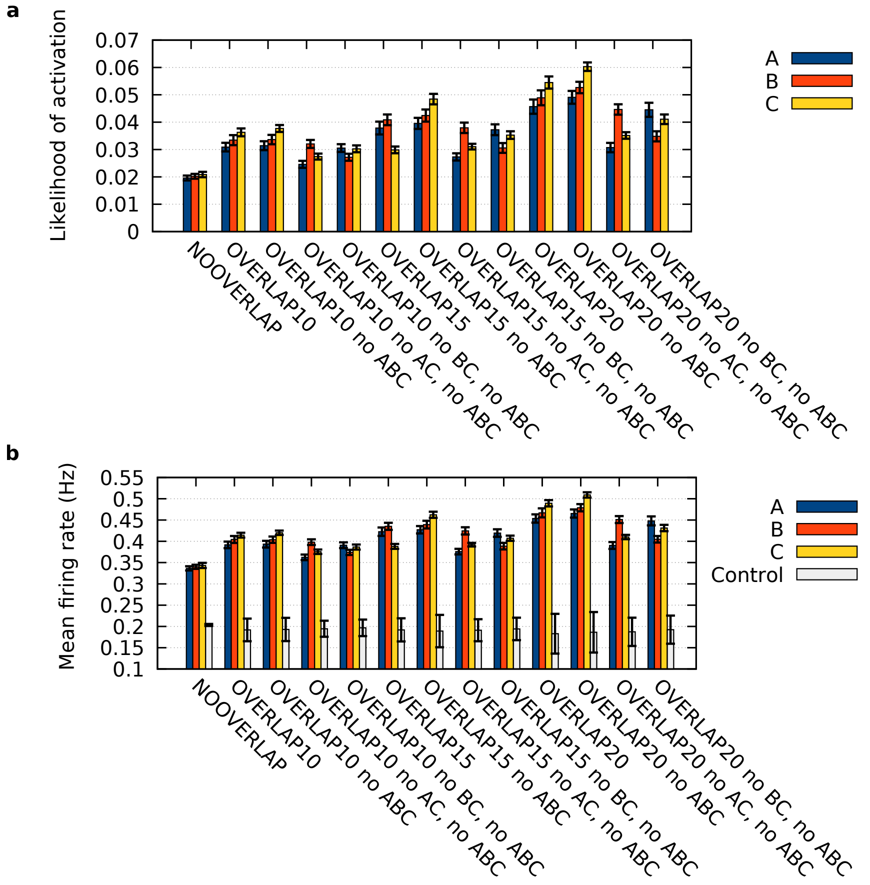


Supplementary Figure S6: Spontaneous activity of long-term memory representations, driven by background noise. Intermediate consolidation protocol (order: $A$-$B$-$C$). **(a)** Overview of likelihood of avalanches across different organizational paradigms, including overlaps of varying size. **(b)** Overview of mean firing rates in the same organizational paradigms as in **a**. Averaged over $10$ trials. Error bars show the 95% confidence interval.

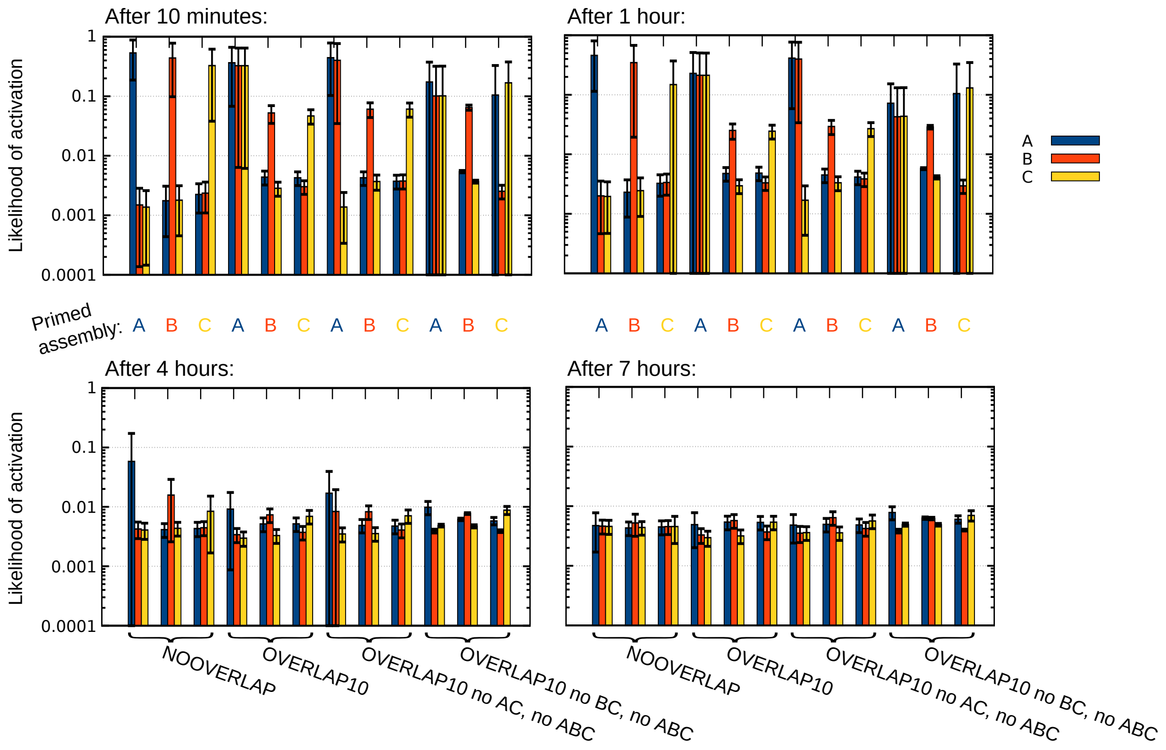


Supplementary Figure S7: Spontaneous activity of long-term memory representations, driven by background noise, after priming stimulus to all neurons of the specified cell assembly. Shown is the likelihood of avalanches across different organizational paradigms at different times after priming. Note the logarithmic scale. Learned with interleaved protocol (order: $A$-$B$-$C$-$A$-$C$-$B$-$B$-$A$-$C$-$A$-$B$-$C$-$C$-$B$-$A$). Averaged over $10$ trials. Error bars show the 95% confidence interval.

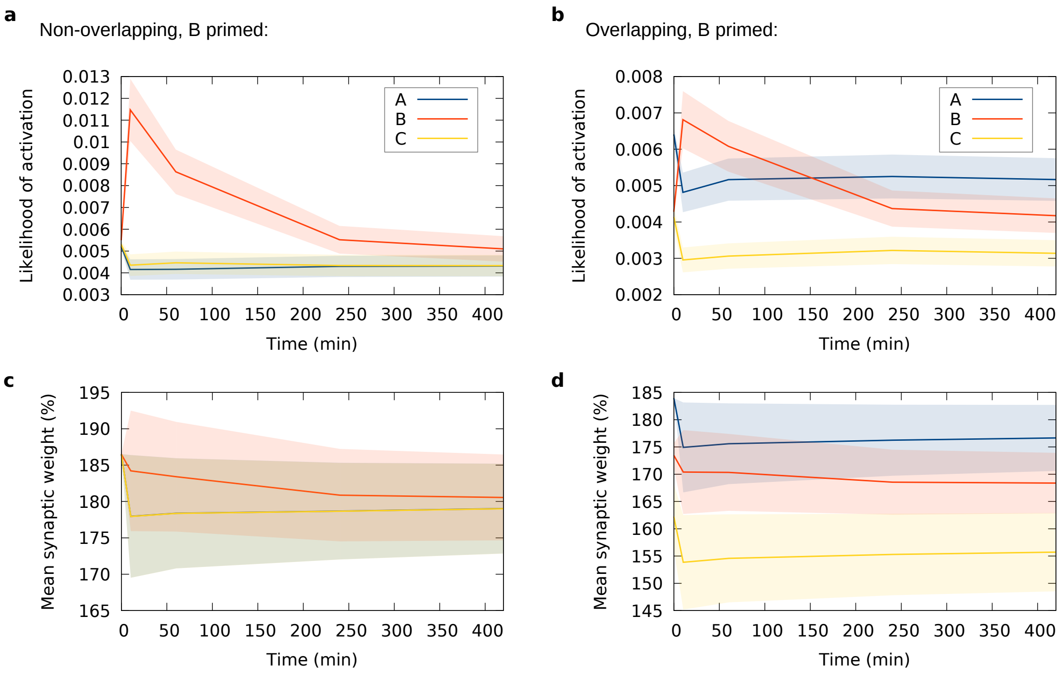


Supplementary Figure S8: Priming on long timescales, enabled by varying degrees of long-term depression, caused by priming stimulation applied to 50% of the neurons of assembly $B$. **(a,b)** Resulting likelihood of avalanches in the three assemblies at different times after priming $B$. The values at time zero show the case before/without priming. NOOVERLAP paradigm in **a**, OVERLAP10 paradigm in **b**. **(c,d)** Temporal development of the mean synaptic weight in the assemblies, for the same cases as in **a** and **b**. Averaged over $10$ trials. Error bands show standard error of the mean. Weight values are relative to the initial value before learning. Learned with interleaved protocol (order: $A$-$B$-$C$-$A$-$C$-$B$-$B$-$A$-$C$-$A$-$B$-$C$-$C$-$B$-$A$).

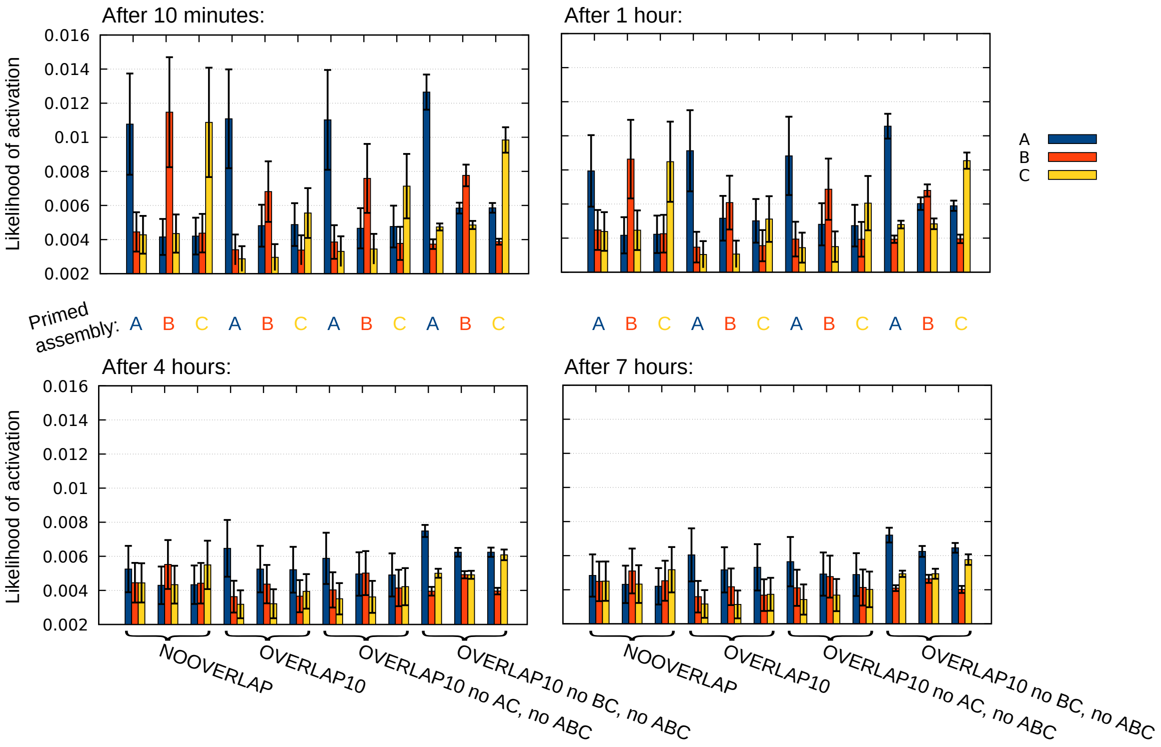


Supplementary Figure S9: Spontaneous activity of long-term memory representations, driven by background noise, after priming stimulus to 50% of the neurons of the specified cell assembly. Shown is the likelihood of avalanches across different organizational paradigms at different times after priming. Learned with interleaved protocol (order: $A$-$B$-$C$-$A$-$C$-$B$-$B$-$A$-$C$-$A$-$B$-$C$-$C$-$B$-$A$). Averaged over $10$ trials. Error bars show the 95% confidence interval.

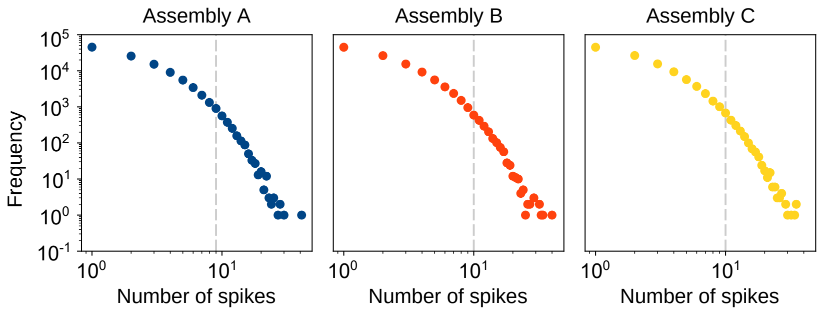


Supplementary Figure S10: Distribution of avalanche size (number of spikes per 10 ms bin), within the three assemblies, after learning and consolidation with standard protocol in the NOOVERLAP paradigm. The dashed lines indicate the border of the 99% quantile for each individual distribution. Data from 10 simulations, each lasting 3 minutes, of the same network.
